# Impact of floral traits on autoluminescence potential in *Phalaenopsis* orchids

**DOI:** 10.64898/2026.04.20.719549

**Authors:** Zhiqing Wang, Jingwei Ma, Junshuo Qiu, Letian Lin, Xiaojuan Wang, Lichuan Chen, Hao Du, Tao Xie, Ruidong Jia, Yuexia Zhang, Jue Ruan, Junxia Wang, Zhimei Li, Peipei Wang

## Abstract

Engineering autoluminescent plants, especially horticultural crops, has recently emerged as a promising research area, with one current approach involving the transgenic introduction of fungal bioluminescence pathway genes. Although autoluminescent plants such as tobacco and petunia have been created, it remains unclear whether all horticultural plants have the potential to be autoluminescent after genetic modification, especially those having flowers with dark colors and thick cuticles. To understand whether, and what if any, floral traits affect autoluminescence potential, we assess the autoluminescence characteristics of 25 representative *Phalaenopsis* cultivars after transient transformation of fungal bioluminescence pathway genes, alongside several key morphological and biochemical traits. Our results demonstrate that autoluminescence characteristics are correlated with floral color lightness, organ textures and epidermal cell types. In contrast, the content of the substrates of luciferin—caffeic acid and tyrosine, and the infiltration ease of inoculation solution into floral organs after injection, have limited effects on autoluminescence characteristics. Autoluminescence intensity can be reasonably predicted using five floral traits investigated, as 80.4% of variation can be explained by these traits. Our study not only identifies specific *Phalaenopsis* cultivars with high potential for developing autoluminescent lines but also provides a selection framework applicable to other horticultural crops.

## Introduction

Autoluminescent organisms are widespread in nature, including numerous marine species, various bacteria, insects like fireflies and beetles, as well as springtails, fungi, and snails^1,2^. This widespread phenomenon contrasts with its rarity in the plant kingdom: no higher plants are autoluminescent, although with ultra-weak luminescence that is far too dim to be perceived by naked eyes and requires particular equipment to be detected^3^. Making flowering plants with autoluminescence has been the target of efforts for over 40 years. Bioluminescent light is normally produced by energy release during the oxidation of luciferin that emits visible light^1^. The luciferins utilized by different organisms are structurally distinct molecules, such as coelenterazine in cnidarians^4^ and FMNH2 (reduced form of flavin mononucleotide) in bacteria^5^. In 1986, Ow *et al*. delivered firefly luciferase into tobacco, but it requires the exogenous application of luciferin (frequently toxic and high-cost) and adenosine triphosphate (ATP) to make the plants produce light, which is normally temporary and non-uniform due to unstable substrate delivery^6^. The bacterial luciferase *Lux* operon (*luxCDABEG*) that composes the light-emission system was delivered into the plastid of tobacco to produce autonomous light without application of luciferin^7^. However, the low brightness, the technical challenges of plastid transformation in plants and potential toxicity of the luciferin substrate for some eukaryotes when nuclearly encoded prevents its widespread applications in plants^8^.

Currently, the only eukaryotic bioluminescence system, which has been well studied and delivered into plants to confer autoluminescence, is from fungi. In fungi, the luciferin is 3-hydroxyhispidin, which is produced by hispidin-3-hydroxylase (H3H) via hydroxylating the hispidin. Luciferase (Luz) adds oxygen to 3-hydroxyhispidin resulting in an unstable high-energy intermediate which emits light subsequently and is decomposed into caffeylpyruvic acid. Caffeylpyruvic acid is then converted to caffeic acid by caffeylpyruvate hydrolase (CPH), while hispidin is synthesized by hispidin synthase (HispS) from caffeic acid^9–12^. Genes encoding *Luz*, *H3H*, *CPH* and *HispS* form a gene cluster in fungi genomes which is relatively conserved in autoluminescent fungi species, with *CPH* sometimes located in other genomic regions ^10,11^. These four genes from *Neonothopanus nambi* were first transformed into tobacco after codon-optimization by Mitiouchkina et al. (2020)^13^.

Khakhar et al. (2020)^8^ added four additional genes into the construction (named as fungal bioluminescence pathway construction, *FBP*): 4’-phosphopantetheinyl transferase (NpgA) from *Aspergillus nidulans* that is required for the post-translational activation of HispS, Rhodobacter capsulatus tyrosine ammonia lyase (RcTAL) and two *Escherichia coli* 4-hydroxyphenylacetate 3-monooxygenase components (HpaB, HpaC); the latter three catalyze caffeic acid synthesis from tyrosine^10^, a molecular that is even more ubiquitous than caffeic acid in higher plants. Shakhova et al. (2024) enhanced the bioluminescence of the *FBP* system by employing a combined strategy of mutagenesis and cross-species comparison to screen for the optimal combination of five key genes: *HispS*, *H3H*, *Luz*, *CPH* and *NpgA*^14^. Zheng et al. (2023) added 4-coumaroyl shikimate/quinate 3’-hydroxylase (C3’H1) from *Brassica napus* to *FBP* (named as enhanced FBP, *eFBP*), leading to enhanced bioluminescence intensity, where C3’H1 facilitates the biosynthesis of caffeic acid and hispidin from p-Coumaroyl shikimate^12^. Sequentially, Ge et al. (2024) integrated a multiplex artificial microRNA (amiR) array into *eFBP* (named as *eFBP2*) to reduce the bypass flow of caffeic acid, leading to 0.7 and 1.0 fold increase in the accumulation of caffeic acid and hispidin, respectively, and further enhanced bioluminescence intensity^15^.

Several plants have been transgenically modified with *FBPs*, including tobacco^8,12,13,16^, hybrid poplar^12^, *Arabidopsis thaliana*^14^, and petunia by the company Light Bio (https://light.bio). These plants tend to have light-colored flowers with thin and soft organs. However, whether dark-colored flowers and those with thick and waxy floral organs have the potential to be autoluminescent after transgenic modification remains unclear. *Phalaenopsis* orchids, having flowers with extraordinarily high diversity in color patterning and organ textures, serve as a good system for assessing the effects of different floral traits on the autoluminescence potential. In addition, *Phalaenopsis* orchids normally have large and gorgeous flowers and exceptionally long flowering phases (2∼3 months), which makes autoluminescent *Phalaenopsis* orchids of inestimable horticultural and economic values.

Here, we assess the autoluminescence characteristics for 25 representative *Phalaenopsis* orchids after transient *eFBP2* transformation, in terms of autoluminescence capability, intensity and duration. These 25 *Phalaenopsis* orchids have flowers with different content of caffeic acid and tyrosine, sizes, color lightness, organ textures, and epidermal cell types. We evaluate the associations between autoluminescence characteristics and these floral traits. Our findings provide insights for the selection of *Phalaenopsis* orchids and other crop plants to develop autoluminescent varieties in the future.

## Methods and materials

### Plant materials and growth conditions

A total of 25 *Phalaenopsis* cultivars were treated in this study (**Fig. S1A, Table S1**). Plants with floral buds and open flowers were purchased from companies in Foshan, where the plants were grown in glass greenhouses with day/night temperatures at 22-30 ℃/17-23 ℃, a natural photoperiod, and light intensities of 220 μmol·m⁻²·s⁻¹.

### Transient transformation of *eFBP2*

The *eFBP2* plasmid was transformed into *Agrobacterium tumefaciens* GV3101 (pSoup) using a freeze-thaw method. Competent cells stored at −80°C were thawed at room temperature until partially melted and subsequently placed on ice. Approximate 100 ng of the *eFBP2* plasmid was added to 100 µL of competent cells, mixed gently, and subjected to sequential inoculation steps: 10 min on ice, 5 min in liquid nitrogen, 5 min in a 37 °C water bath, and 5 min on ice. Subsequently, 900 µL of antibiotic-free lysogeny broth (LB) medium was added, and the cells were incubated at 28 °C with shaking at 220 rpm for 2 h to allow recovery. After centrifugation at 6,000 rpm for 1 min, the supernatant was largely discarded, and the pellet was resuspended before being plated onto LB agar plates containing 50 µg/mL kanamycin and 25 µg/mL rifampicin. The plates were incubated inverted at 28 °C in the dark for 2 d. Single colonies were then inoculated into 3 mL LB medium supplemented with the same antibiotics and cultured at 28 °C, with shaking at 220 rpm for 2 d. Plasmid presence was confirmed by PCR using *eFBP2*-*Luz* primers (forward: 5′-CTAGAATTCATGAGAATCAACATCTCACT-3′; reverse: 5′-TTCGTCGACTTACTTAGCGTTCTCAACGA-3′) with 1 µL of bacterial culture as the template. Verified *Agrobacterium* cultures were expanded and stored at −80°C for subsequent experiments.

To initiate the transient transformation, *Agrobacterium* cultures carrying *eFBP2* vectors were grown in 150 mL LB medium with the same antibiotics as above at 28 °C with shaking at 230 rpm until the OD600 reached 1.8 (approximately 14 h). The cells were harvested by centrifugation at 6,000 rpm for 8 min at room temperature, resuspended in MMA buffer (10 mM MES, 10 mM MgCl₂, 100 µM acetosyringone, pH 5.6) to an OD600 of 0.8, and then incubated in the dark for 2 h. Silwet L-77 was added to a final concentration of 0.005% (v/v), and the suspension was mixed thoroughly. Prior to infiltration, plants were withheld from watering for several days to facilitate a degree of solute concentration within tissues. The bacterial suspension was injected into the abaxial surface of tepals using a 1-mL syringe equipped with a needle. Injection was performed until complete tissue infiltration was observed, with care taken to prevent repeated puncturing to minimize tissue damage. A minimum of four plants per cultivar were utilized, and all the open flowers and flowers prior to anthesis on these plants were injected. Subsequently, plants were covered with black plastic bags to preserve humidity and maintained in the dark at room temperatures for ∼24 h (∼30 °C in the summer, while ∼22 °C in the autumn), then were transferred to a growth room (∼22 °C on average, 220 µmol·m⁻²·s⁻¹ LED illumination, 14 h light/10 h dark), and irrigation was withheld for a further 2–3 d.

### Assessment of autoluminescence characteristics

The autoluminescence characteristics of *eFBP2*-transformed plants were first recorded two days after injection (DAI) photographically, and were subsequently monitored at 12-h (in the autumn experiments) or 24-h (in the summer experiments) intervals until the autoluminescence disappeared completely. For each *Phalaenopsis* cultivar, only the flower with the highest autoluminescence intensity was photographed with a Canon EOS 90D camera (ISO 25600, the minimum aperture ranging from f/3.5 to f/5.6, 30-second exposure in Bulb mode) in the dark field. To grade cultivars with different autoluminescence intensities, the image with the highest brightness was first converted into grayscale image. Three brightest regions (5.4 mm × 7.0 mm in size) were selected, and their grayscale values were determined using ImageJ. The averaged grayscales per image was used to represent the autoluminescence intensity of the cultivar in target. The autoluminescence intensity can be categorized into four grades: none (grayscale = 0), low (0 < grayscale ≤ 30), medium (30 < grayscale ≤ 90), or high (grayscale > 90). Autoluminescence duration was defined as the number of days during which autoluminescence was detectable, commencing 24 h after transfer to the growth room and ending when it was no longer visible to the camera and/or the naked eye.

### High Performance Liquid Chromatography analysis

To measure the content of caffeic acid and tyrosine in flowers of *Phalaenopsis* cultivars, the petal samples were ground in liquid nitrogen and thoroughly mixed. Exactly 0.1 g of powder was weighed and placed in a 15 mL centrifuge tube. For caffeic acid, 3 mL of methanol was added into the 15 mL centrifuge tube, and the mixture was ultrasonicated at 4 ℃ for 1 h, and then centrifuged at 4000 rpm for 10 min. A suitable amount of the supernatant was filtered through a 0.22 μm filter for testing. The samples were analyzed using a High Performance Liquid Chromatography (HPLC) system (Agilent 1100) with C18 column (4.6 mm ✕ 250 mm, 5 μm, Agilent). The HPLC analysis was performed using an isocratic elution with a mobile phase of acetonitrile and 0.1% phosphoric acid in water (40:60, v/v). The flow rate was set at 1.0 mL/min, and the column temperature was maintained at 30 °C, and the detection wavelength was 300 nm.

For tyrosine, 5 mL of 6 M HCl was added into the 15 mL centrifuge tube (containing petal powder), and was mixed thoroughly on a shaker for 5 min. The sample was transferred to a preheated oven at 105 °C for hydrolysis for 24 h. After hydrolysis, the sample was cooled immediately with cold water to room temperature, and checked whether HCl should be supplemented to make up for the deficiency. The sample was neutralized by adding 5 mL of 6 M NaOH and vortexing vigorously for 5 min, and then was centrifuged at 4000 rpm for 10 min. A mixture of 0.5 mL of the supernatant, 0.5 mL of bicarbonate buffer (pH 9.0), and 0.5 mL of 2,4-dinitrofluorobenzene (DNFB) was transferred into a 15 mL brown centrifuge tube, and incubated in a light-protected water bath at 60.0 ± 0.5 °C for 60 min, and was cooled immediately to room temperature with cold water. The mixture was added 3.5 mL of phosphate buffer (pH 7.0), mixed thoroughly and incubated at room temperature in the dark for 15 min. The supernatant was filtered through a 0.22 μm membrane filter, and was analyzed using a HPLC system (Agilent 1100) with C18 column (4.6 mm ✕ 250 mm, 5 μm, Agilent). The HPLC analysis was conducted with a binary gradient mobile phase composed of solvent A (acetonitrile and methanol in a 90:10 ratio) and solvent B (0.02 M monobasic sodium phosphate and sodium hydrogen phosphate). The gradient started from 14% (v/v) solvent A and 86% (v/v) solvent B, then the concentration of solvent A was increased to 70% (v/v) over 44 min, and then reset to 14% (v/v) at 44.01 min for column re-equilibration. The method utilized a constant flow rate at 1.0 mL/min, and a column temperature of 38 °C, and UV detection at 360 nm.

Caffeic acid and tyrosine were identified by comparing their retention times (18 and 46 min, respectively) with those of authentic standards (Shanghai Yuanye Bio-Technology Co., Ltd) and quantified using external calibration curves generated from serially diluted standard solutions.

### Measurement of floral traits

Floral traits recorded for 25 *Phalaenopsis* cultivars included flower size, color lightness, texture of floral organs, epidermal cell type and ease of infiltration (**Table S2**). Specifically, the diameter of a mature flower (typically the first flower at the base of the inflorescence) was measured per plant, with five biological replicates per cultivar. Floral color lightness was quantified using the CIELAB color space, where the L represents perceptual lightness, which ranges from 0 (black) to 100 (white). Three representative points were sampled on the petal per cultivar, and the averaged L value was recorded. For *P*. ‘Michelle’, the L value was measured for the lip instead of the petal since the lip was the only organ with autoluminescence. The floral color lightness was categorized into three grades: dark (L ≤ 50), medium (50 < L < 70) and light (L ≥ 70).

Texture of floral organs was classified as waxy or velvety according to established criteria in refs^17,18^, where waxy tepals are more brittle than velvety ones. To simplify the analysis, we gently folded the petals and classified cultivars with broken petals as waxy while the ones with unbroken petals as velvety.

## Results

### Experiment design and the measure of autoluminescence characteristics

A total of 25 cultivars (**Fig. S1A**) that differed in several floral key traits—including size, color lightness, venation-associated stripe, spot or patch pattern, and floral organ texture—were treated with *eFBP2*. Two batches of experiments were performed (**Table S1**): one in the autumn of 2025 with room temperature about 22 °C (22 cultivars, **Fig. S2**), the other in the summer of 2025 when the room temperature was about 30 °C (all 25 cultivars, **Fig. S3**). Jointly considering the ease of injection operation and the endogenous biochemical activity of flowers at different development stages, only flowers that were immediately prior to anthesis (S2), partially open (S3), fully open (S4) and those began to wilt (S5) were injected with *eFBP2*-containing bacterial suspension, while those at bud stage (S1) were not used (**Fig. S1B,C**). Three to six flowers were treated per plant and were numbered according to their positions on the inflorescence (**Fig. S1C**). At least 14 flowers from at least 4 plants were treated per cultivar for the summer experiments, while ≥ 3 flowers from ≥ 1 plant were treated per cultivar for the autumn experiments (**Table S1**).

After transient transformation with *eFBP2*, flowers of certain cultivars displayed clear green autoluminescence at two days after injection (DAI) (**Fig. 1A-C**, **Fig. S2,S3**). Dark-field photos of flowers were recorded using Bulb mode of the camera, with a consistent 30-second exposure at ISO 25600 (**Methods**). Flowers were photographed every 12 h (**Fig. S2**) or 24 h (**Fig. S3**) till the bioluminescence light was no longer detectable, and the duration of autoluminescence was recorded. To ensure comparability of autoluminescence characteristics across cultivars, the characteristics of the flower with the highest autoluminescence intensity were recorded for each cultivar and each experimental batch. This strategy was adopted because we noticed that within each cultivar, the flowers capable of autoluminescence exhibited relatively comparable autoluminescence properties in the same experimental batch (**Fig. S4**). Cultivars were further divided into four categories according to their autoluminescence intensity, which was calculated as the averaged grayscale across three brightest regions on their flowers (**Fig. 1D, Methods**). Specifically, cultivars with high intensity had grayscale > 90, medium intensity with grayscale between 30 and 90, low intensity with grayscale ≤ 30, while those with no autoluminescence had grayscale = 0 (**Fig. 1E**, **Table S1**). For flowers capable of autoluminescence, the highest light intensity was observed between 2 and 4 DAI, and later on, the light intensity decreased gradually (**Fig. 1C, Fig. S2,S3**). The duration of autoluminescence ranged from 1 to 8 days (**Fig. 1F**, **Fig. S5A**). Cultivars with higher autoluminescence intensity tended to have longer duration of the autoluminescence (**Fig. 1G, Fig. S5B**).

**Fig. 1.**
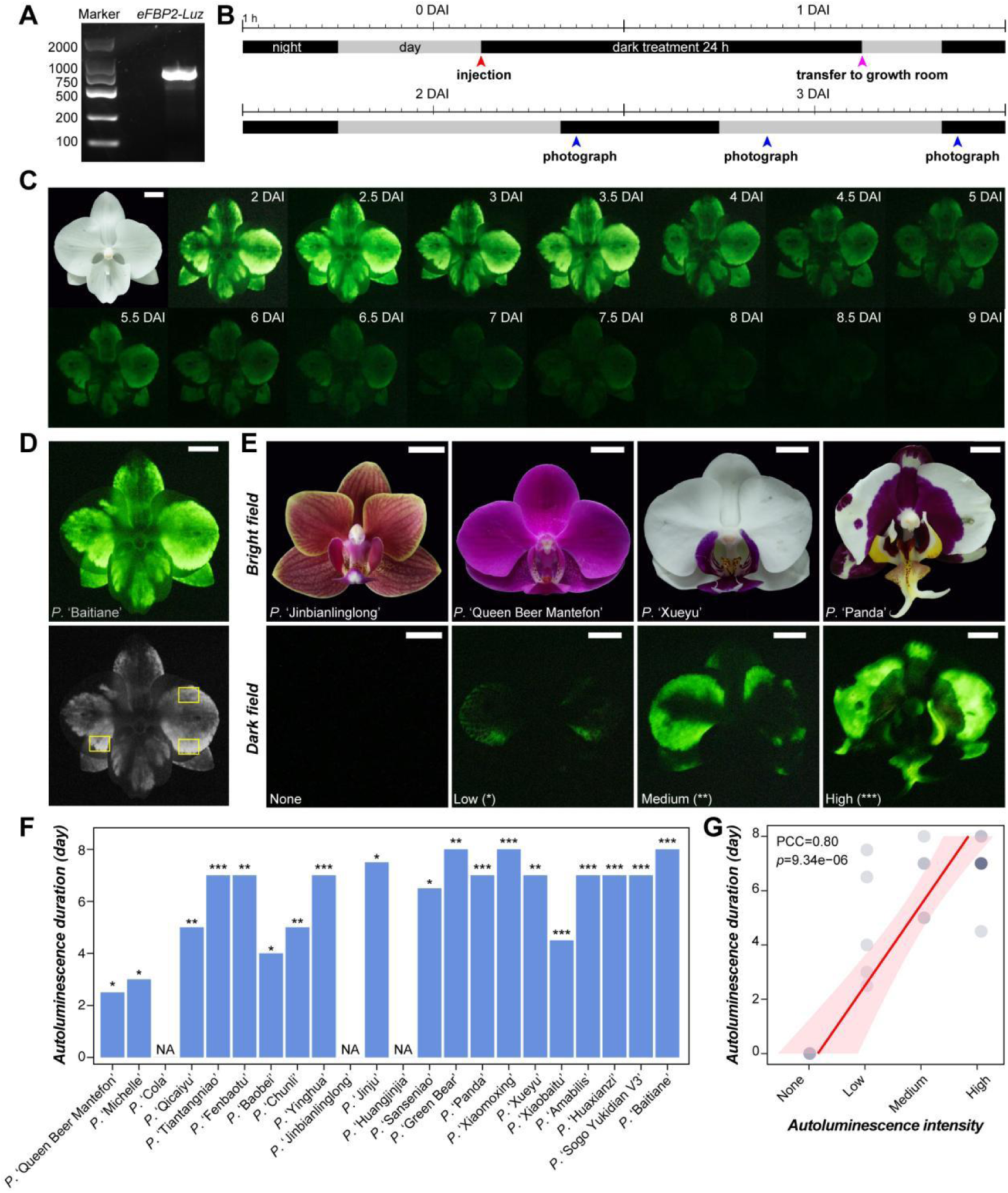
Measure of autoluminescence characteristics. (**A**) Gel electrophoresis image showing the PCR production of *eFBP2*-*Luz*. (**B**) Schematic timeline illustrating the schedule of the injection of *eFBP2*-containing inoculation solution, dark treatment, transfer to growth room, and photography. DAI: day after injection. (**C**) Bright-field image of *P*. ‘Baitiane’, and dark-field images taken every 12 h after *eFBP2* transient transformation. (**D**) The dark-field image of *P*. ‘Baitiane’ and its corresponding grayscale image. Three yellow rectangles indicate the brightest regions measured for the grayscale values. (**E**) Bright- and dark-field images of four cultivars, representing the ones with no, low, medium and high autoluminescence, respectively. (**F**) Autoluminescence duration of 22 *Phalaenopsis* cultivars. Asterisk above each bar denotes the autoluminescence intensity of the corresponding cultivar: low (*), medium (**), and high (***). NA: not applicable. (**G**) Correlation between autoluminescence intensity and duration. PCC: Pearson correlation coefficient. Scale bar: 1 cm.

We found substantial effects of environmental temperatures on all the three autoluminescence characteristics measured: autoluminescence capability, intensity, and duration. Among 22 cultivars examined in the autumn experiments, 9, 5 and 5 showed high, medium and low bioluminescence intensity, respectively, while 3 failed to emit autoluminescence (**Fig. 1F, Fig. S2**). On the other hand, among 25 cultivars examined in the autumn experiments, the corresponding numbers were 4, 6, 7 and 8, respectively (**Fig. S3, Fig. S5A**). *P*. ‘Cola’, *P*. ‘Jinbianlinglong’, and *P*. ‘Huangjinjia’ were not autoluminescent regardless of room temperatures. When comparing the autoluminescence characteristics of 22 overlapping cultivars, the autumn experiments had more cultivars with autoluminescence, higher intensity, and longer duration than the summer experiments (**Fig. S6**). Particularly, *P*. ‘Baitiane’ failed to emit autoluminescence in all the summer experiments but had high intensity in the autumn ones; *P*. ‘Michelle’ and *P*. ‘Sanseniao’ were not autoluminescent in the summer experiments, but had low intensity in the autumn experiments. These findings underscore the need for optimizing experiments to identify suitable plant materials for genetic modification with *FBPs*. Nevertheless, the autoluminescence characteristics of these cultivars between two batches of experiments are positively correlated (Cramér’s V ≥ 0.57, *p* of Chi-square test ≤ 3.52e-03, **Fig. S6**). Unless otherwise specified, we focused on the results of autumn experiments to facilitate the following analyses.

### The content of caffeic acid and tyrosine had limited effects on autoluminescence characteristics

Considering that the *eFBP2* system used caffeic acid as the substrates to synthesize hispidin, and integrated the *Rhodotorula glutinis* TAL and HpaB, HpaC to convert tyrosine to caffeic acid^15^, we asked whether the autoluminescence characteristics are mainly determined by the content of caffeic acid and tyrosine in floral organs. Given that flowers that were partially open or newly open (S3) tended to have higher autoluminescence intensity (**Fig. 2A**, **Table S1**), we measured the content of these two compounds in petals at S3 for six representative cultivars (**Fig. 2B,C**, **Methods**). For *P.* ‘Amabilis’, petals at S4 and S5 were also examined to assess the dynamic fluctuation of tyrosine and caffeic acid content during flower development.

**Fig. 2.**
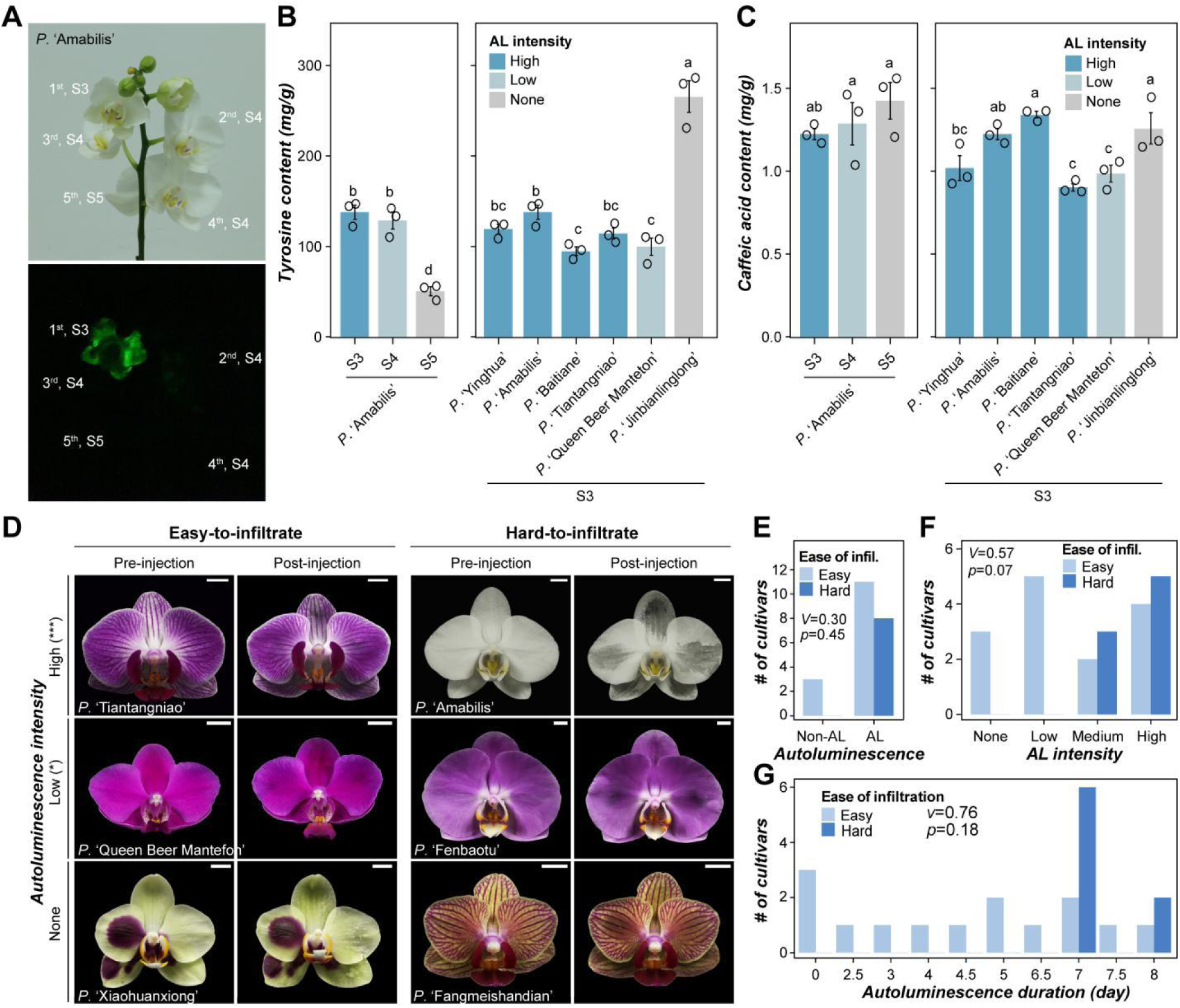
Limited influences of caffeic acid and tyrosine content and the ease of infiltration on autoluminescence characteristics. (**A**) Bright-field (upper) and corresponding dark-field (lower) images of *P*. ‘Amabilis’ inflorescence, illustrating the position-dependent autoluminescence after *eFBP2* transformation. (**B,C**) The tyrosine and caffeic acid content in petals of six cultivars at stages S3, S4 and S5. Error bars: ± standard deviation (n = 3). Different lowercase letters indicate significant differences between groups (ANOVA, *p* < 0.05). (**D**) Images of six cultivars—with no (None), low (*) or high (***) autoluminescence intensity—before and after being injected with *eFBP2*-containing inoculation solution, showing differences in the ease of liquid infiltration into floral tissues. Results of the remaining 19 cultivars were shown in **Fig. S7**. Scale bar: 1 cm. (**E-G**) Correlation between autoluminescence capability (**E**), intensity (**F**) and duration (**G**) and the ease of infiltration. AL: autoluminescence; *V*: Cramér’s V; *p*: *p*-value of Chi-square test.

Surprisingly, the content of these two compounds was not correlated with autoluminescence characteristics (**Fig. 2B,C**): the non-autoluminescent cultivar *P*. ‘Jinbianlinglong’ had higher content of tyrosine than all the other four cultivars with high autoluminescence intensity; the content of caffeic acid in *P*. ‘Jinbianlinglong’ was also higher than or comparable with those in other cultivars. In addition, although the tyrosine content decreased along flower development in *P.* ‘Amabilis’ (**Fig. 2B**), the caffeic acid content increased gradually (**Fig. 2C**). This is potentially because *eFBP2* rewires the tyrosine → caffeic acid → hispidin → luciferin metabolic flux, and the caffeic acid and tyrosine contents in wild-type flowers do not represent the metabolite pool available for the hispidin and luciferin synthesis during transient transformation. These results suggest that other floral traits, rather than the substrate content in floral organs, determine the autoluminescence characteristics of *Phalaenopsis* orchids after *eFBP2* transformation.

### Relationships between autoluminescence characteristics and the ease of liquid infiltration into floral organs

During the experiments, we noticed that floral organs of different cultivars differed in the ease of infiltration with the inoculation solution. In 16 cultivars, the inoculation solution spread readily and evenly throughout the tissues after injection (‘easy-to-infiltrate’), while in the other 9 cultivars, the inoculation solution tended to remain localized near the injection site (‘hard-to-infiltrate’) (**Fig. 2D**, **Fig. S7**). The variation in ease of infiltration reflected the differences in cell composition and arrangement in these floral organs, which might influence the diffusion of *eFBP2*-containing bacteria in these organs as well. In addition, to ensure a valid comparison, more injection attempts were performed for these hard-to-infiltrate cultivars to achieve a relatively uniform distribution of inoculation solution, which may diminish the vitality of floral organs and lead to a reduction in autoluminescence potential consequently. We detected strong (Cramér’s V=0.57 and 0.76 for autoluminescence intensity and duration, respectively), but not significant (*p*-value = 0.07 and 0.18), correlations between the ease of infiltration and autoluminescence characteristics (**Fig. 2E-G**). Surprisingly, the eight hard-to-infiltrate cultivars tended to show medium or high autoluminescence intensity lasting 7–8 days, whereas the remaining 14 easy-to-infiltrate cultivars exhibited more even distribution across intensity and duration levels. This is contrary to our initial hypothesis—harder to infiltrate, more injections, lower vitality and lower autoluminescence potential. These results indicate that the observed variation in autoluminescence characteristics did not mainly stem from treatment differences.

### Correlation between autoluminescence characteristics and other floral traits

#### Flower size

Among 25 cultivars, 3, 12 and 10 had large (diameter > 10 cm), medium (7 cm ≤ diameter ≤ 10 cm) and small (< 7 cm) flowers, respectively (**Fig. S8**, **Table S2,S3**). We identified weak to moderate but not significant correlations (Cramér’s V < 0.65, *p* ≥ 0.44) between autoluminescence characteristics and flower sizes (**Fig. 3A-D**) for 22 cultivars examined in the autumn experiments. All the three cultivars with large flowers had medium or high autoluminescence intensity lasting seven days. Flowers with medium sizes also tended to have higher intensity and longer duration, while those with small sizes exhibited relatively even distribution across intensity and duration levels. A similar tendency was also observed in the summer experiments (**Fig. S5F-H**).

**Fig. 3.**
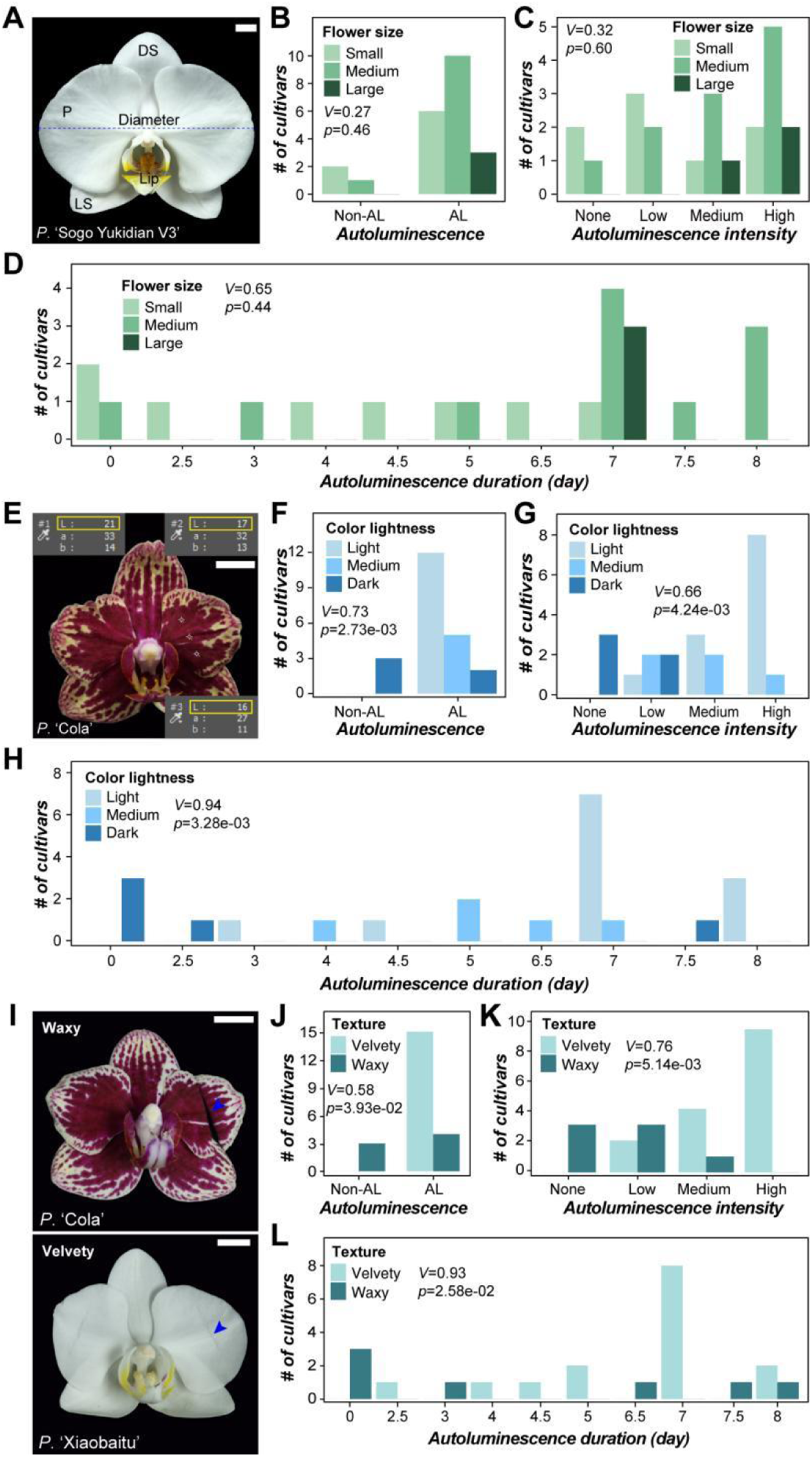
Correlation between autoluminescence characteristics and three floral traits. (**A**) Image of *P*. ‘Sogo Yukidian V3’ showing how the flower diameter (flower size) is measured. DS: dorsal sepal; P: petal; LS: lateral sepal. (**B-D**) Correlation between the autoluminescence characteristics and flower sizes. (**E**) Image of *P*. ‘Cola’ showing the measure of color lightness (L) of three representative regions on its petals. (**F-H**) Correlation between the autoluminescence characteristics and color lightness of petals. (I) Images of *P.* ‘Cola’ and *P.* ‘Xiaobaitu’ after gently folding, illustrating cultivars with waxy and velvety petals, respectively. (**J-L**) Correlation between the autoluminescence characteristics and texture of floral organs. AL: autoluminescence; *V*: Cramér’s V; *p*: *p*-value of Chi-square test. Scale bar: 1 cm.

#### Floral color lightness

Considering the extraordinary complexity of color patterning in *Phalaenopsis* orchids (**Fig. S1A**), to facilitate the analysis, we measured the color lightness of these flowers (**Fig. 3E, Methods**) instead of the color *per se*. Flowers with light colors (average Lab values ≥ 70) tended to be autoluminescent, with higher intensity and longer durations (Cramér’s V ≥ 0.66, *p* ≤ 4.24e-03, **Fig. 3F-H**). Particularly, in flowers with venation-associated stripes—such as those of *P*. ‘Yinghua’, *P*. ‘Tiantangniao’, and *P*. ‘Qicaiyu’, and flowers with patches—such as *P*. ‘Panda’, the dark colored stripes and patches tended to have much lower autoluminescence intensity than their neighboring regions (**Fig. S2**). In flowers of *P*. ‘Fenbaotu’, only the base of sepals and petals, and the whiskers had detectable autoluminescence, where the colors were lighter than other regions (**Fig. S2**). In addition, the white sepals and petals of *P*. ‘Xueyu’ were autoluminescent while the violet lip was not; the white lip of *P*. ‘Michelle’, rather than the dark red sepals and petals, had detectable autoluminescence. These results suggest that dark colors may obstruct the emission of autoluminescence—a possibility that requires cytological and spectral absorbance data to confirm—and that color lightness of the floral organs is a good indicator of autoluminescence potential after genetic modification. There were also exceptions, such as flowers of *P*. ‘Queen Beer Mantefon’, which were dark purple across the whole flower but had detectable, albeit weak, autoluminescence, indicating that other factors should also contribute to the autoluminescence characteristics.

#### Floral organ texture

Floral organs in *Phalaenopsis* orchids can be roughly classified into waxy and velvety ones according to their textures^17^. Waxy organs are normally thicker and brittler, covered with thicker cuticles, compared with velvety ones^17,19^. To simplify the analysis, we classified these 25 cultivars into waxy and velvety based solely on the tendency of their petals to break when gently folded (**Fig. 3I**, **Table S2**). We found strong and significant correlations (Cramér’s V ≥ 0.58, *p* ≤ 2.58e-02) between all three autoluminescence characteristics and the floral organ texture (**Fig. 3J-L**). All the 15 cultivars with velvety floral organs were autoluminescent after *eFBP2* transformation, 9 of which had high intensity (**Fig. 3J,K**). In contrast, seven cultivars with waxy floral organs were not autoluminescent (three) or had low intensity (four). Similar correlation was also observed for the summer experiments (**Fig. S5L-N**). These results suggest that the thick cuticle on the surface of floral organs may also hinder the emission of autoluminescence, which should be taken into account in the selection of suitable cultivars.

#### Epidermal cell types on floral organs

We examined the freehand sections of four types of perianth—dorsal and lateral sepals, petals, and the lip, and photographed the epidermal cell types in the center region of each organ (**Fig. 4A**). Given that only the autoluminescence characteristics on the adaxial surfaces of floral organs were recorded, we only examined the epidermal cells on the adaxial surfaces as well. *Phalaenopsis* cultivars examined in this study displayed extraordinary diversity in epidermal cell types (**Fig. 4B**, **Fig. S9**). These cell types can be classified into several categories according to their shapes and height^20^: flat, domelike with different heights, conical with different shapes, and papillate. Different cell types may coexist in some cultivars, such as domelike and conical cells on the petals of *P*. ‘Qicaiyu’ (**Fig. 4B**). In addition, cells of these cultivars displayed a somewhat continuous range of shapes, making it difficult to classify them into distinct types, particularly for domelike cells. Jointly considering these concerns, we further divided the cells into four categories according to their height/width (h/w) ratios (averaged across three representative cells): type Ⅰ (h/w = 0), corresponding to the flat cell type; type Ⅱ (0 < h/w ≤ 0.5), including the flat domelike cells and some of the conical cells with pyramid-like shape; type Ⅲ (0.5 < h/w ≤ 1.0), the most prevalent type on sepals and petals among *Phalaenopsis* orchids; type Ⅳ (h/w > 1.0), including partial elongated domelike, conical and papillate cells (**Fig. 4B, Fig. S9, Table S4**). Type Ⅰ cells were never observed on sepals and petals, while type Ⅳ cells were absent from the lips (**Fig. 4C**).

**Fig. 4.**
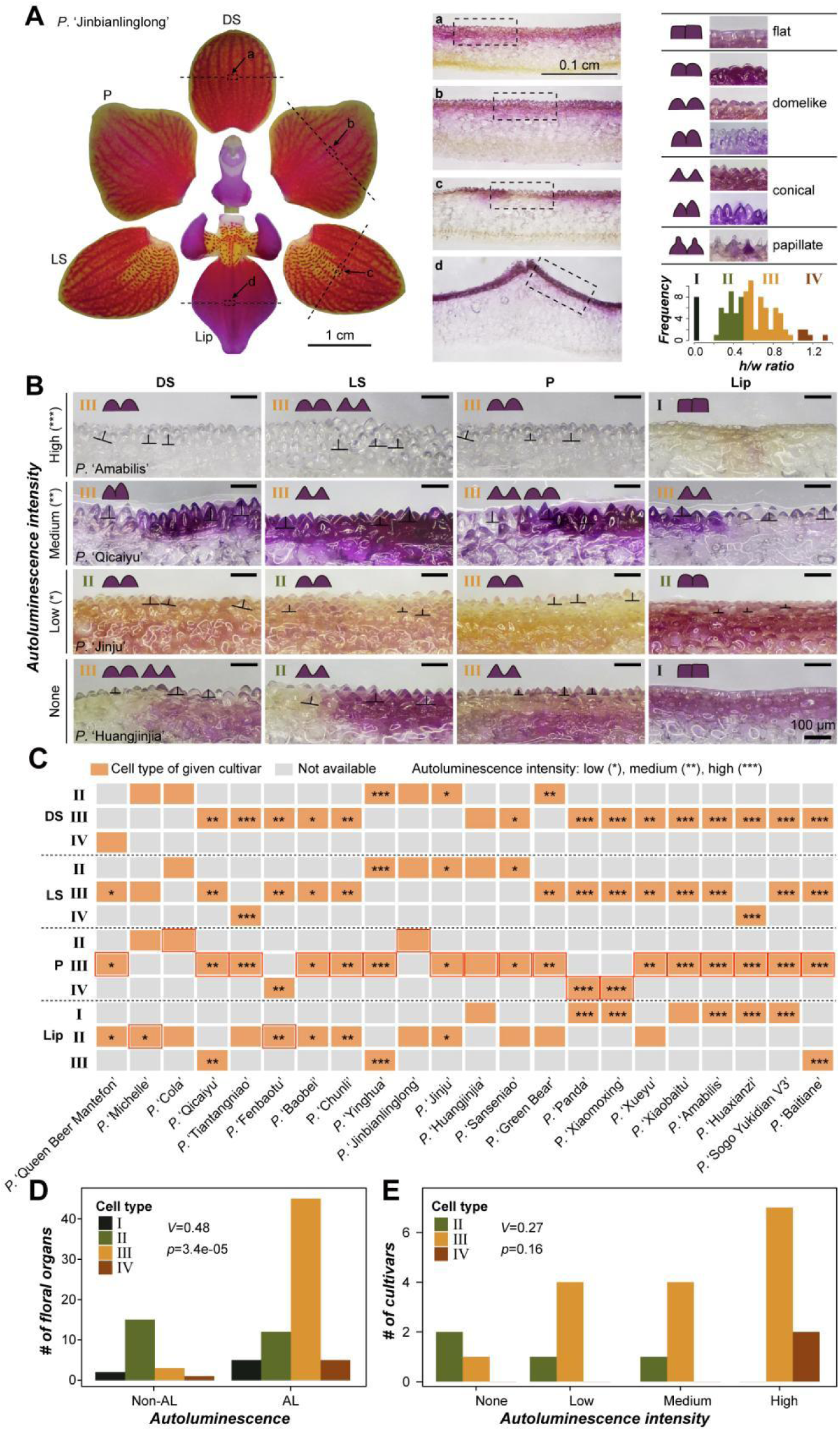
Correlation between autoluminescence characteristics and epidermal cell types of floral organs. (**A**) *P*. ‘Jinbianlinglong’ floral organ images showing positions of transverse sections (dash lines) and the locations of epidermal cells examined (a-d). Scale bar: 1 cm for the organ images and 0.1 cm for the transverse section images. (**B**) Images of floral organ epidermal cells from four cultivars exhibiting no (None), low (*), medium (**) and high (***) autoluminescence intensity, respectively. Seven epidermal cell types are illustrated above (**B**). The classification of cells into four categories (types I–IV) based on their average height/width ratios (n = 3) was shown as well. Scale bar: 100 μm. Results of the remaining 21 cultivars were shown in **Fig. S9**. (**C**) Distribution of epidermal cell types on different floral organs of 22 cultivars. Results of the remaining three cultivars were shown in **Fig. S5O**. Orange rectangle: the cell type of the floral organ in a given cultivar. Rectangles with *, **, and *** indicate organs with low, medium and high autoluminescence intensity, respectively. DS, dorsal sepal; LS, lateral sepal; P, petal. A red rectangle indicates the floral organ with the highest autoluminescence intensity per flower, or the petal if the cultivar was not autoluminescent. (**D**) Correlation between autoluminescence capability and epidermal cell types in all floral organs. (**E**) Correlation between autoluminescence intensity and cell types in the representative floral organ with the highest intensity per cultivar. *V*: Cramér’s V; *p*: *p*-value of Chi-square test.

We found moderate but significant correlation (Cramér’s V = 0.48, *p* = 3.4e-06) between cell types and autoluminescence capability (**Fig. 4D**). Floral organs whose epidermal cells exhibited high h/w ratios tended to show autoluminescence following *eFBP2* transformation (**Fig. 4C**). In contrast, organs with flattened epidermal cells did not. The only one floral organ with type Ⅳ but having no autoluminescence was the dorsal sepal of *P*. ‘Queen Beer Mantefon’, which had dark purple floral organs. Petals of this cultivar showed weak autoluminescence, and only restricted regions on the lateral sepals and the lip were autoluminescent.

Since different floral organs of a single flower exhibited distinct autoluminescence intensity and cell types (**Fig. S2, S3, S7**), the association analysis between cell type and autoluminescence intensity was performed only on the floral organ with the highest autoluminescence intensity per cultivar (red rectangles in **Fig. 4C**). No significant correlation (Cramér’s V = 0.27, *p* = 0.16) was identified (**Fig. 4E**). Nevertheless, two cultivars with type Ⅳ epidermal cells had high autoluminescence intensity on their petals, and cultivars with type Ⅲ cells also tended to have higher intensity than those with type Ⅱ cells.

### Prediction of autoluminescence characteristics from floral traits via multivariate modeling

When showing different floral traits and autoluminescence characteristics of 22 cultivars within a single heatmap (**Fig. 5A**), we found all three autoluminescence characteristics were clustered with floral color lightness and organ texture, consistent with what we have found above. Three non-autoluminescent cultivars, namely *P*. ‘Huangjinjia’, *P*. ‘Cola’ and *P*. ‘Jinbianlinglong’, shared common floral traits: waxy textures, dark colors, type Ⅱ or Ⅲ epidermal cells, small or medium size and easy-to-infiltrate. On the contrary, cultivars with high or medium autoluminescence intensity tended to have light-colored, velvety, medium-sized or large flowers, with epidermal cells exhibiting relatively high height/width (h/w) ratios. Then, we asked if the autoluminescence potential of a cultivar after transformed with *FBPs* can be predicted jointly from all the floral traits examined here. Considering the relatively small sample size (n = 22), we simply built regression models using the lm function in R ^21^ to estimate the contribution of each floral trait to the autoluminescence characteristics, rather than established machine learning models to predict the autoluminescence characteristics in a training/test split scheme. The variance inflation factors (VIFs) for all the five floral traits were smaller than 2.5 (**Fig. 5B**), indicating that multicollinearity among the five floral traits was negligible.

**Fig. 5.**
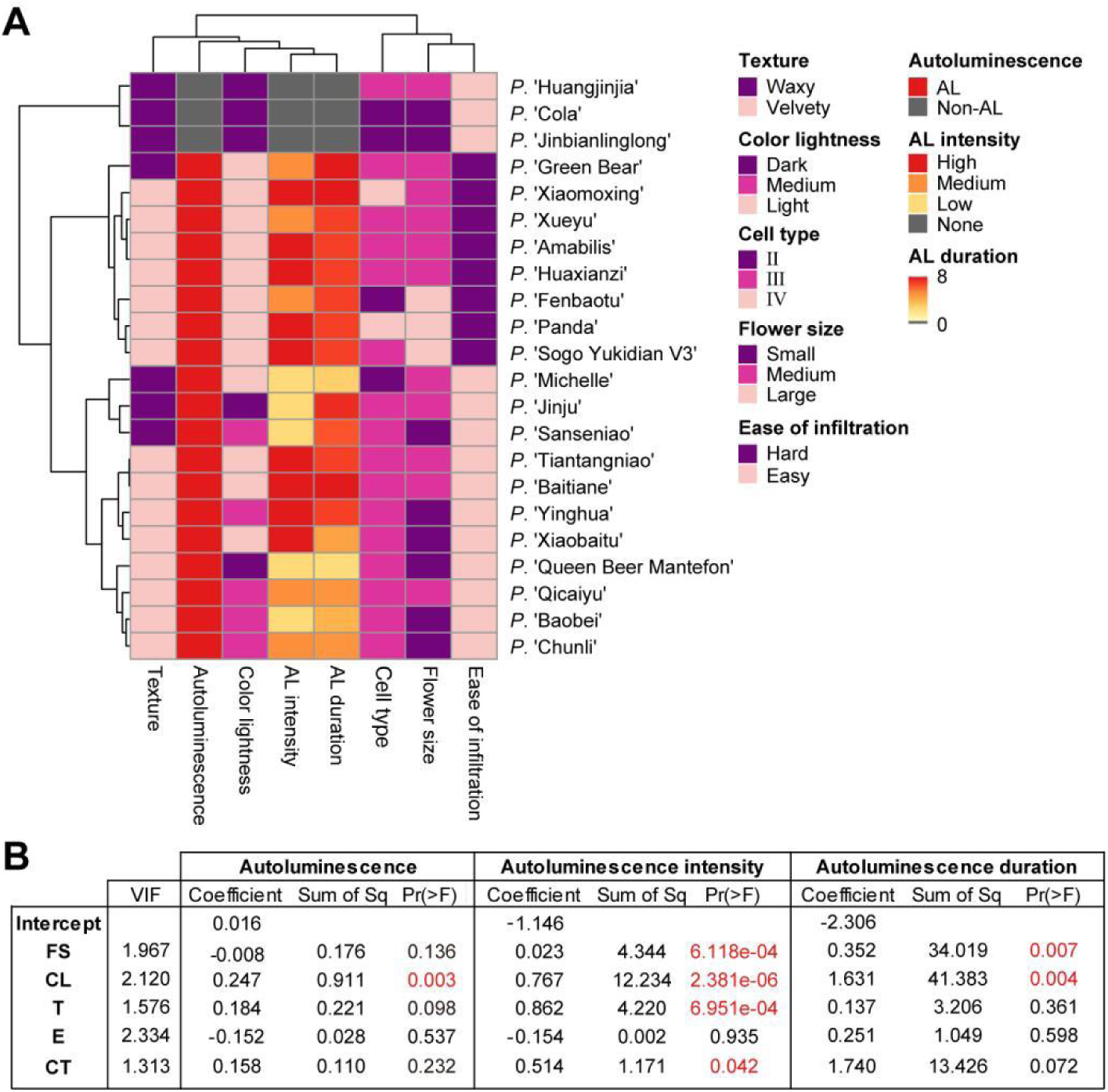
Prediction of autoluminescence characteristics using floral traits. (**A**) Clustering heatmap among autoluminescence characteristics and floral traits of 22 *Phalaenopsis* cultivars. Color scale: corresponding categories or values of different autoluminescence characteristics and floral traits, as illustrated in the legend. (**B**) Regression models for three autoluminescence characteristics. Sum of Sq: the additional explained variation in autoluminescence characteristics when a given floral trait was added into the regression models. AL: autoluminescence; FS: flower size; CL: color lightness; T: texture of floral organs; E: ease of infiltration; CT: cell type on the representative floral organ, as indicated in Fig. 4C; VIF: variance inflation factor; Sq: Squares; Pr(>F): Probability (greater than F), namely *p*-value.

For the capability of autoluminescence after *eFBP2* transformation, the regression model was: AL = 0.016 - 0.008FS + 0.247CL + 0.184T - 0.152E + 0.158CT (**Fig. 5B**), where AL: autoluminescence; FS: flower size; CL: color lightness; T: texture of floral organs; E: ease of infiltration; CT: cell type on the representative floral organ. The adjusted R^2^ of this model was 0.421 (F-statistic = 4.047, *p* = 0.014), indicating 42.1% of variation in autoluminescence capability can be explained by variation in these floral traits across cultivars. Among these traits, only floral color lightness had a significant contribution to the prediction. For the autoluminescence intensity, more variation can be explained using floral traits (adjusted R^2^ = 0.804, F-statistic = 18.27, *p* = 4.164e-06). Flower size, color lightness, organ texture and cell type on the representative floral organ contributed significantly to the prediction. For the autoluminescence duration, 49.6% of the variation was explained using floral traits (F-statistic = 5.135, *p* = 5.343e-03). Flower size and color lightness contributed significantly to the prediction. These results demonstrate that color lightness consistently contributed to the prediction of three autoluminescence characteristics, while other floral traits contributed to predictions of one or two characteristics, and infiltration ease contributed to none of the predictions.

Models for the summer experiments led to poorer predictions for all three autoluminescence characteristics (adjusted R^2^ = 37.4%, 62.4% and 37.0% for autoluminescence capability, intensity and duration, respectively), potentially due to the influences of higher temperatures. The contributions of floral traits to predictions in summer experiments were similar to those in autumn experiments, except that flower size and organ texture contributed significantly to predictions of all three autoluminescence characteristics, while color lightness did not contribute to the prediction of autoluminescence capability in the summer experiments (**Fig. S10**). Taken together, all the three autoluminescence characteristics, especially autoluminescence intensity, can be reasonably predicted or explained by these five floral traits.

## Discussion

### Influences of environmental temperatures on autoluminescence characteristics

By assessing the autoluminescence characteristics of 25 *Phalaenopsis* cultivars after transient *eFBP2* transformation in two batches of experiments, we demonstrated the influences of environmental temperatures on autoluminescence potential, as higher temperatures decreased autoluminescence capability, intensity and duration. These temperature-sensitive autoluminescence characteristics were also reported in other studies^10,12^. Zheng *et al*. (2023) found that the autoluminescence intensity decreased rapidly upon high temperature treatment although the caffeic acid amount increased^12^. Kotlobay *et al*. (2018) demonstrated that the luciferase loses its activity at temperatures above 30 ℃^10^. In addition, high temperatures above 30 ℃ reduced the efficiency of T-DNA transfer to plants due to the thermosensitivity of *Agrobacterium tumefaciens*^22–24^. Nevertheless, the association between autoluminescence characteristics and floral traits was similar across two experimental batches conducted under different environmental temperatures, suggesting that the observed tendency in our study is relatively robust.

### Association between floral traits and autoluminescence characteristics

*Phalaenopsis* flowers with light colors tended to be capable of autoluminescence, have high autoluminescence intensity and long duration. In flowers with dark-colored stripes or patches, these stripes or patches showed significantly lower autoluminescence intensity than their neighbouring regions. This association between color lightness and autoluminescence intensity was also found in other plants. For example, in autoluminescent tobacco, the light-colored flowers normally have higher autoluminescence intensity than the green stem and leaves, although all the cells of the whole plant carry the *FBPs*^16^. Furthermore, we had never observed autoluminescence from *Phalaenopsis* leaves after transient *eFBP2* transformation regardless of cultivars, potentially due to their dark green colors and cuticles on the surface^25^. Flower size also contributed to the prediction of autoluminescence intensity and duration. Cultivars with large flowers are mainly progenies of *P. amabilis* and *P. aphrodite*^26^, and large flowers in *Phalaenopsis* are normally associated with light colors and velvety textures. Flower size can serve as an indicator for selecting potential materials for modification with *FBPs*, but this indicator might not be suitable for other horticultural crops. Floral organ texture and epidermal cell type contributed to the prediction of autoluminescence intensity in both experimental batches, suggesting that these two traits should be taken into account in the selection of candidate materials as well.

### Limited effects of infiltration ease on autoluminescence potential

On the contrary, infiltration ease had the least contributions to the predictions of autoluminescence characteristics among other floral traits. The infiltration ease was not used as a proxy of *Agrobacterium* infection efficiency, since the physical fluid spread does not capture T-DNA delivery, bacterial attachment, or host compatibility. Instead, the ease of infiltration affected flower treatment: hard-to-infiltrate flowers required more injection attempts to achieve a relatively uniform distribution of the inoculation solution. Different treatments may therefore affect floral organ vitality and, consequently, autoluminescence potential. The absence of a significant association between autoluminescence characteristics and infiltration ease suggests that the different autoluminescence characteristics we recorded for these *Phalaenopsis* orchids are unlikely to result from treatment fluctuation or non-uniform delivery of the inoculation solution across floral tissues.

### Methodological implications

It is worth noting that the screening results from transient *eFBP2* transformation in this study might not directly predict the suitability for stable autoluminescent line development, due to several potential reasons. First, during transient *eFBP2* transformation, not all cells of the floral organs are capable of expressing *eFBP2* and producing luciferin, whereas every single cell in a stable autoluminescent line—including epidermis cells—is expected to have that capability. Second, autoluminescent characteristics in transient transformation are affected by multiple factors, such as flower stage and condition, experimental temperature, *Agrobacterium* infection efficiency, substrate availability, and potentially other unrecognized factors. Finally, *eFBP2* rewires the tyrosine → caffeic acid → hispidin → luciferin metabolic flux, which is thought to differ significantly between transiently transformed wild-type flowers and stable transgenic flowers. This might explain why no correlation was identified between autoluminescence characteristics and caffeic acid/tyrosine content in this study. A more robust and generalizable conclusion could be refined after stable transgenic experiments on representative cultivars.

## Conclusion

In this study, we demonstrated that not all flowers possess the potential for autoluminescence after transient *eFBP2* transformation, and that flowers of different *Phalaenopsis* cultivars differ in autoluminescence capability, intensity and duration. Furthermore, owing to the considerably high diversity and complexity in color patterning and epidermal cell types in *Phalaenopsis* orchids, different flowers on a single plant, different organs within a single flower, and even different regions of a single floral organ exhibit distinct autoluminescence characteristics. Generally, floral organs with light colors, velvety textures and raised epidermal cells show high autoluminescence potential. Our findings shed light on the selection of potential materials for the time- and labor-consuming transgenic experiments required to create autoluminescent horticultural plants.

## Author contributions

P.W., Z.L. and J.W. conceived and designed this study. H.D provided the eFBP2 construct; Z.W., J.M., Z.L., L.L. and J.Q. performed the transient transformation experiments with help from L.C. and X.W.; Z.W., J.M and Z.L. measured the flower diameters and assessed the floral organ textures; Z.W. and J.M. took the bright- and dark-field images; Z.W., Z.L., L.L. and J.Q. collected flower images before and after injection; Z.W., Z.L., J.M, J.Q., and L.L. collected the autoluminescence data; J.M wrote the scripts for statistics analysis; P.W., Z.W., J.M., Z.L., L.L. and J.Q. wrote the manuscript with inputs from all authors. All authors read and approved the final manuscript.

## Data availability statement

All the raw data used in this study have been submitted in the supplementary information.

## Conflict of interests

The authors declare that they have no conflict of interests.

## Acknowledgement

This work was supported by the National Natural Science Foundation of China (32370241), the collaborative project (HXQS2024009 from Prof. Jue Ruan) to P.W.; the Innovation Program of Chinese Academy of Agricultural Sciences to J.R.; the Natural Science Foundation of Guangdong Province, China (2026A1515011083) to J.W.; National Key R&D Program of China (2023YFD1600505) to R.J.

## Supplementary figure legends

**Fig. S1 Flowers of 25 *Phalaenopsis* cultivars and the definition of flowering stages.** (**A**) Flowers of 25 *Phalaenopsis* cultivars. (**B**) Different flower stages of *P.* ‘Yinghua’. S1, bud stage; S2, immediate pre-anthesis; S3, partially open; S4, fully opened; S5, initial wilting. (**C**) Inflorescence of *P.* ‘Fenbaotu’. Flowers are numbered sequentially according to their positions on the inflorescence, where 1^st^ refers to the topmost flower that is injectable (at S2, S3 or S4). Scale bar: 1 cm.

**Fig. S2 Dark-field images of 19 cultivars that were autoluminescent after transient *eFBP2* transformation in the autumn experiments.** A total of 22 cultivars were treated. Dark-field images were taken every 12 h till the autoluminescence was not detectable. Images captured at the last DAI had their brightness and contrast adjusted to facilitate the illustration of weak autoluminescence. AL: autoluminescence; DAI: day after injection. Scale bar: 1 cm.

**Fig. S3 Dark-field images of 20 cultivars that were autoluminescent after transient *eFBP2* transformation in the summer or autumn experiments.** A total of 25 cultivars were treated. Images of three cultivars that were not autoluminescent in the summer experiments but autoluminescent in the autumn experiments were also shown. Dark-field images were taken every 24 h till the autoluminescence was not detectable. Images captured at the last DAI had their brightness and contrast adjusted to facilitate the illustration of weak autoluminescence. The corresponding dark-field images from the autumn experiments showing autoluminescence are displayed on the right. AL: autoluminescence; DAI: day after injection. Scale bar: 1 cm.

**Fig. S4 Correlation of autoluminescence characteristics between two autoluminescent flowers per cultivar in the autumn experiments.** Red rectangles indicate the results used in the main analyses. Dark-field images were taken every 12 h till the autoluminescence was not detectable. Images captured at the last DAI had their brightness and contrast adjusted to facilitate the illustration of the weak autoluminescence. AL: autoluminescence; DAI: day after injection. Scale bar: 1 cm.

**Fig. S5 Results of the summer experiments.** (**A**) Autoluminescence duration of 25 *Phalaenopsis* cultivars. Asterisk above each bar denotes the autoluminescence intensity of the corresponding cultivar: low (*), medium (**), and high (***). NA: not applicable. (**B**) Correlation between autoluminescence intensity and duration. PCC: Pearson correlation coefficient. (**C-N**) Correlation between the autoluminescence capability (**C,F,I,L**), intensity (**D,G,J,M**) and duration (**E,H,K,N**) and the ease of infiltration (**C-E**), flower size (**F-H**), color lightness (**I-K**), and organ texture (**L-N**). (**O**) Distribution of epidermal cell types on different floral organs of three cultivars. Results of the remaining 22 cultivars were shown in **Fig. 4C**. Orange rectangle: the cell type of the floral organ in a given cultivar. Rectangle with * indicates the organ with low autoluminescence intensity. DS, dorsal sepal; LS, lateral sepal; P, petal. A red rectangle indicates the floral organ with the highest autoluminescence intensity per flower, or the petal if the cultivar was not autoluminescent. (**P**) Correlation between autoluminescence capability and epidermal cell types in all floral organs. (**Q**) Correlation between autoluminescence intensity and cell types in the representative floral organ with the highest intensity per cultivar. AL: autoluminescence; *V*: Cramér’s V; *p*: *p*-value of Chi-square test.

**Fig. S6 Correlation of autoluminescence intensity (A) and duration (B) between two batches of experiments.** *V*: Cramér’s V; *p*: *p*-value of Chi-square test.

**Fig. S7 The ease of liquid infiltration into floral organs of 19 cultivars.** Images of flowers before and after being injected with *eFBP2*-containing inoculation solution. Results of the remaining six cultivars were shown in **Fig. 2D**. Scale bar: 1 cm.

**Fig. S8 Flower diameters of 25 cultivars.** Error bars: ± standard deviation (n=5).

**Fig. S9 Epidermal cell types of floral organs in 21 cultivars.** Seven types of epidermal cells were illustrated on the right. The classification of cells into four categories (types Ⅰ, Ⅱ, Ⅲ and Ⅳ) according to their average height/width ratios (n = 3) was shown as well. Results of the remaining four cultivars were shown in **Fig. 4B**. Scale bar: 100 μm.

**Fig. S10 Prediction of autoluminescence characteristics using floral traits for the summer experiments.** (**A**) Clustering heatmap among autoluminescence characteristics and floral traits of 25 *Phalaenopsis* cultivars. Color scale: corresponding categories or values of different autoluminescence characteristics and floral traits, as illustrated in the legend. (**B**) Regression models for three autoluminescence characteristics. Sum of Sq: the additional explained variation in autoluminescence characteristics when a given floral trait was added into the regression models. AL: autoluminescence; FS: flower size; CL: color lightness; T: texture of floral organs; E: ease of infiltration; CT: cell type on the representative floral organ, as indicated in **Fig. 4C** and **Fig. S5O**; VIF: variance inflation factor; Sq: Squares; Pr(>F): Probability (greater than F), namely *p*-value.

## Supplementary tables

**Table S1** Autoluminescent data statistics of 25 *Phalaenopsis* cultivars

**Table S2** Floral traits of 25 *Phalaenopsis* cultivars

**Table S3** Flower diameters of 25 *Phalaenopsis* cultivars

**Table S4** Classification of epidermal cells of 25 *Phalaenopsis* cultivars

## Notes

### Competing Interest Statement

The authors have declared no competing interest.

### Summary of Updates

Have updated the writting of this manuscrpt.

